# Large-scale annotated dataset for cochlear hair cell detection and classification

**DOI:** 10.1101/2023.08.30.553559

**Authors:** Christopher J. Buswinka, David B. Rosenberg, Rubina G. Simikyan, Richard T. Osgood, Katharine Fernandez, Hidetomi Nitta, Yushi Hayashi, Leslie W. Liberman, Emily Nguyen, Erdem Yildiz, Jinkyung Kim, Amandine Jarysta, Justine Renauld, Ella Wesson, Punam Thapa, Pierrick Bordiga, Noah McMurtry, Juan Llamas, Siân R. Kitcher, Ana I. López-Porras, Runjia Cui, Ghazaleh Behnammanesh, Jonathan E. Bird, Angela Ballesteros, A. Catalina Vélez-Ortega, Albert SB Edge, Michael R. Deans, Ksenia Gnedeva, Brikha R. Shrestha, Uri Manor, Bo Zhao, Anthony J. Ricci, Basile Tarchini, Martin Basch, Ruben S. Stepanyan, Lukas D. Landegger, Mark Rutherford, M. Charles Liberman, Bradley J. Walters, Corné J. Kros, Guy P. Richardson, Lisa L. Cunningham, Artur A. Indzhykulian

**Affiliations:** Eaton Peabody Laboratories, Mass Eye and Ear, Boston, MA, 02114, USA; Department of Otolaryngology, Head and Neck Surgery, Harvard Medical School, Boston, MA, 02114, USA; Speech and Hearing Biosciences and Technology graduate program, Harvard University, Cambridge, MA, 02138, USA; Section on Sensory Cell Biology, National Institute on Deafness and Other Communication Disorders, National Institutes of Health, Bethesda, MD, 20814, USA; Department of Otolaryngology, Head and Neck Surgery, Vienna General Hospital and Medical University of Vienna, 1090 Vienna, Austria; Department of Otolaryngology, Washington University School of Medicine, St. Louis, MO, USA; Department of Neuroscience, Washington University School of Medicine, St. Louis, MO, USA; The Jackson Laboratory, Bar Harbor, ME, 04609, USA; Department of Otolaryngology-Head and Neck Surgery, Case Western Reserve University School of Medicine, Cleveland, OH, 44106, USA; The University of Mississippi Medical Center, Dept. of Otolaryngology – Head and Neck Surgery, Jackson, MS, USA; Department of Neurobiology, Spencer Fox Eccles School of Medicine at the University of Utah, Salt Lake City, UT 84112, USA; Department of Otolaryngology – Head & Neck Surgery, Spencer Fox Eccles School of Medicine at the University of Utah, Salt Lake City, UT, 84132, USA; Eli and Edythe Broad CIRM Center for Regenerative Medicine and Stem Cell Research, University of Southern California, Los Angeles, CA, 90033, USA; Tina and Rick Caruso Department of Otolaryngology-Head and Neck Surgery, University of Southern California, Los Angeles, CA, 90033, USA; Waitt Advanced Biophotonics Center, Salk Institute for Biological Studies, 10010 N. Torrey Pines Road, La Jolla, CA, 92037, USA; Department of Cell and Developmental Biology, University of California San Diego, La Jolla, CA, 92093; Department of Otolaryngology-Head and Neck Surgery, Indiana University School of Medicine, Indianapolis, IN, 46202, USA; Department of Otolaryngology, Stanford University School of Medicine, Stanford, CA, 94305, USA; Department of Molecular and Cellular Physiology, Stanford University School of Medicine, Stanford, CA, 94305, USA; Department of Medicine, Tufts University, Boston, 02111, MA, USA; Graduate School of Biomedical Science and Engineering (GSBSE), University of Maine, Orono, ME, 04469, USA; Department of Neurosciences, Case Western Reserve University School of Medicine, Cleveland, OH, 44106, USA; Department of Otolaryngology, Washington University, 660 S. Euclid Avenue, Campus Box 8115, St. Louis, MO, 63110, USA; Sussex Neuroscience, School of Life Sciences, University of Sussex, Brighton, United Kingdom; Department of Physiology, University of Kentucky, Lexington, KY, 40536, USA; Section on Sensory Physiology and Biophysics, National Institute on Deafness and Other Communication Disorders, National Institutes of Health, Bethesda, MD, 20814, USA; Department of Pharmacology and Therapeutics, University of Florida, Gainesville, FL, 32610, USA; Myology Institute, University of Florida, Gainesville, FL, 32610, USA

**Keywords:** hair cells, outer hair cell, inner hair cell, cochlea, detection, annotation, machine-learning-ready data

## Abstract

Our sense of hearing is mediated by cochlear hair cells, localized within the sensory epithelium called the organ of Corti. There are two types of hair cells in the cochlea, which are organized in one row of inner hair cells and three rows of outer hair cells. Each cochlea contains a few thousands of hair cells, and their survival is essential for our perception of sound because they are terminally differentiated and do not regenerate after insult. It is often desirable in hearing research to quantify the number of hair cells within cochlear samples, in both pathological conditions, and in response to treatment. However, the sheer number of cells along the cochlea makes manual quantification impractical. Machine learning can be used to overcome this challenge by automating the quantification process but requires a vast and diverse dataset for effective training. In this study, we present a large collection of annotated cochlear hair-cell datasets, labeled with commonly used hair-cell markers and imaged using various fluorescence microscopy techniques. The collection includes samples from mouse, human, pig and guinea pig cochlear tissue, from normal conditions and following *in-vivo* and *in-vitro* ototoxic drug application. The dataset includes over 90,000 hair cells, all of which have been manually identified and annotated as one of two cell types: inner hair cells and outer hair cells. This dataset is the result of a collaborative effort from multiple laboratories and has been carefully curated to represent a variety of imaging techniques. With suggested usage parameters and a well-described annotation procedure, this collection can facilitate the development of generalizable cochlear hair cell detection models or serve as a starting point for fine-tuning models for other analysis tasks. By providing this dataset, we aim to supply other groups within the hearing research community with the opportunity to develop their own tools with which to analyze cochlear imaging data more fully, accurately, and with greater ease.

## BACKGROUND / SUMMARY

The cochlea is the hearing organ of the mammalian inner ear. It contains thousands of highly specialized mechanically sensitive cells, called hair cells. In response to the mechanical force of sound, vibrations of the tympanic membrane are transmitted along the cochlea in a travelling wave, leading to the stimulation of sensory hair cells. Organized along the length of the cochlea in an exquisitely stereotyped sensory epithelium within the structure of the organ of Corti, hair cells are subdivided into two types: three rows of outer hair cells (OHC) which amplify mechanical vibration and one row of inner hair cells (IHC) which translate that vibration into neural signals^1^. The cochlea is tonotopically organized^2^, with the critical frequency to which hair cells most strongly respond determined by location. Hair cells at the basal end of the cochlea respond most strongly to high frequency sounds while those at the apex respond to low frequencies. Hair cells are essential for sound perception, however they are sensitive to damage or loss as a result of cochlear insult^3^. Mammalian cochlear hair cells are terminally differentiated and do not regenerate^4,5^. As such, hair cell death leads to permanent sensorineural hearing loss.

The severity of damage, and critically, where it has occurred, are required to fully characterize the extent of any cochlear trauma. Similarly, efforts to develop otoprotective therapeutics require detailed quantification of hair cell survival. Typically, hair cells at the base of the cochlea are considered most vulnerable to damage, putting the detection of high frequency sounds at the highest risk of loss, from both insult and aging. Similarly, tonotopic effects are also often evaluated when testing various therapeutics and insults to the cochlea. A common route of drug delivery to the cochlea is through the round window membrane at the base of the cochlea^6^. This creates a gradient of therapeutic concentration and potentially, efficacy^7^. Therefore, when evaluating trauma and/or treatment outcomes, researchers often need to assess hair cell loss on a large scale, along the tonotopic axis. However, with thousands of hair cells per cochlea this task becomes prohibitive and is rarely performed, necessitating the development and use of automated analysis tools. Finally, although it is often sufficient to quantify each subtype of hair cell and record their position along the length of the cochlea, more sophisticated assessment and quantification might reveal additional characteristics beyond cell loss, such as hair bundle disarray^8^. To facilitate various types of analyses, it is desirable for detection tools to determine the area each individual cell occupies, rather than just their presence.

Object detection is a well-studied branch of computer vision^9^ and machine learning algorithms have been successfully implemented to detect instances of hair cells, classify their type (OHC vs IHC), and to denote their area by a bounding box^10-12^. The most successful object detection algorithms rely on deep neural networks and are often trained on a large set of examples to maximize their accuracy. This either requires the manual generation of very large annotated datasets, or the use of a model pretrained with a large, general dataset and “fine-tuning” it on a smaller subset of specialized training data^13^. Models, trained on data for a variety of cell types are increasingly accessible, however, they may fail when presented with new or atypical data^14^. Hair cells are such a case and are not widely represented in any public dataset to date. Therefore, any deep learning model to detect hair cells requires *de novo* generation of a hair-cell-specific dataset. Comprehensive datasets of annotated cochlear hair cells are challenging to create as the cochlea is studied in different animal models, with a wide range of pathological conditions, and at different developmental time points, all of which can dramatically affect hair cell appearance. Hair cells may be imaged with different light microscopy techniques, staining procedures, labeling procedures, or sample preparation procedures. Furthermore, tissue morphology complexities are compounded when studying developmental alterations or cochlear insults resulting in unique patterns of histopathological findings^15^. The lack of available comprehensive annotated dataset hinders the development of cochlear analysis tools.

To meet this need, we have compiled and annotated an open source, deep-learning-ready cochlear hair cell detection dataset. This collection of data is a large collaborative effort: cochlear samples were prepared, and imaging datasets collected by independent hearing research laboratories, curated to be representative of techniques across the field of auditory neuroscience (summarized in **Table 1**). Each image was annotated by experts (**Figure 1**), and carefully validated to ensure accuracy. We further provide recommendations and exemplar code for data pre-processing, strategies for proper train-test splitting, sampling procedures for whole cochlear datasets, and processes necessary for the fine tuning of future deep learning models. This dataset is the foundational dataset for the Hair Cell Analysis Toolbox, a deep learning-based software for cochlear analysis we previously reported^10^. It has therefore been demonstrated as sufficient to train accurate detection models. It is our hope that this dataset can be used by other research groups to train and benchmark automated hair cell detection and classification algorithms, and thereby aid in their future development.

**Table 1:**
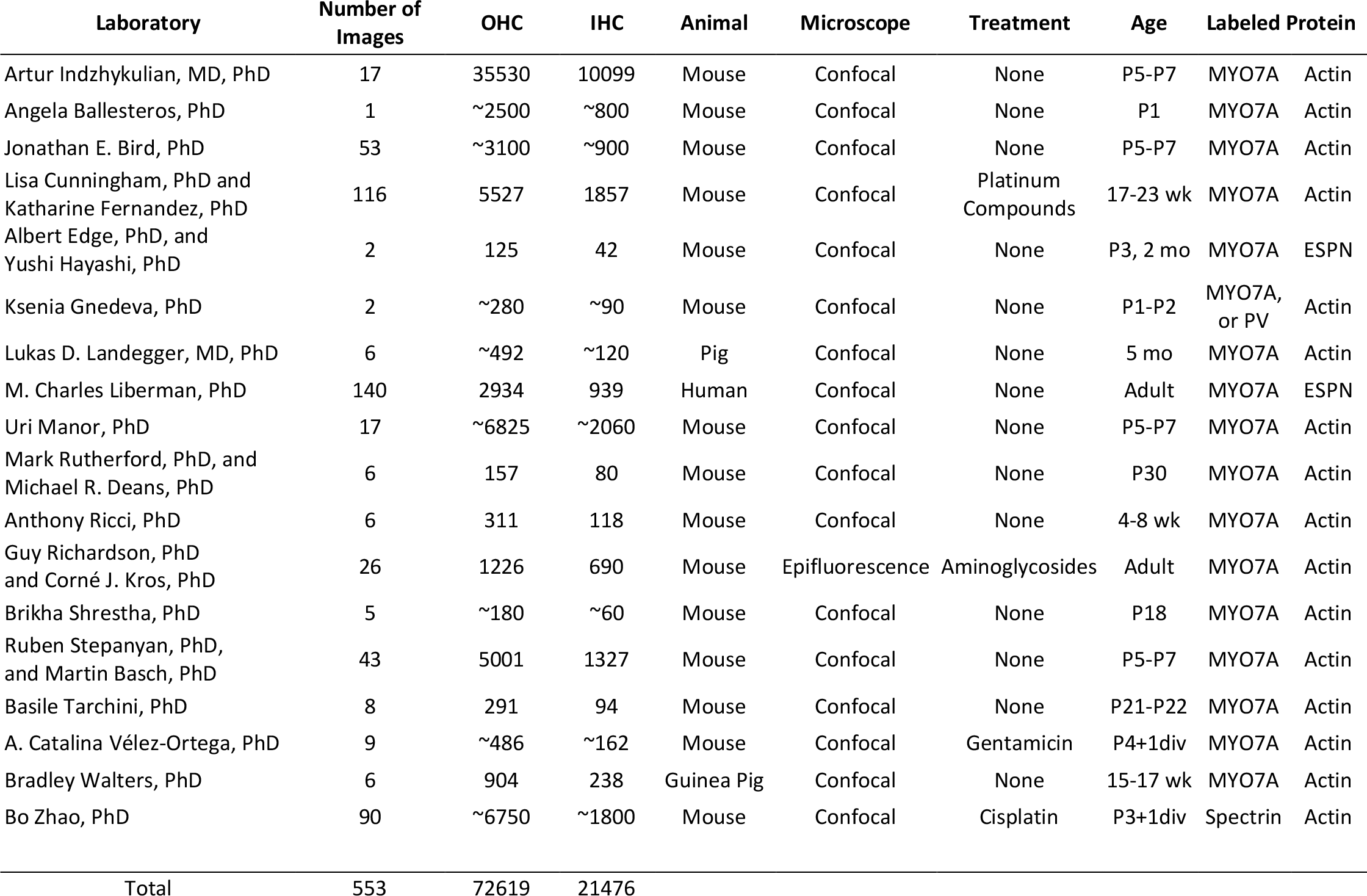
Summary of Annotated Data.

**Figure 1:**
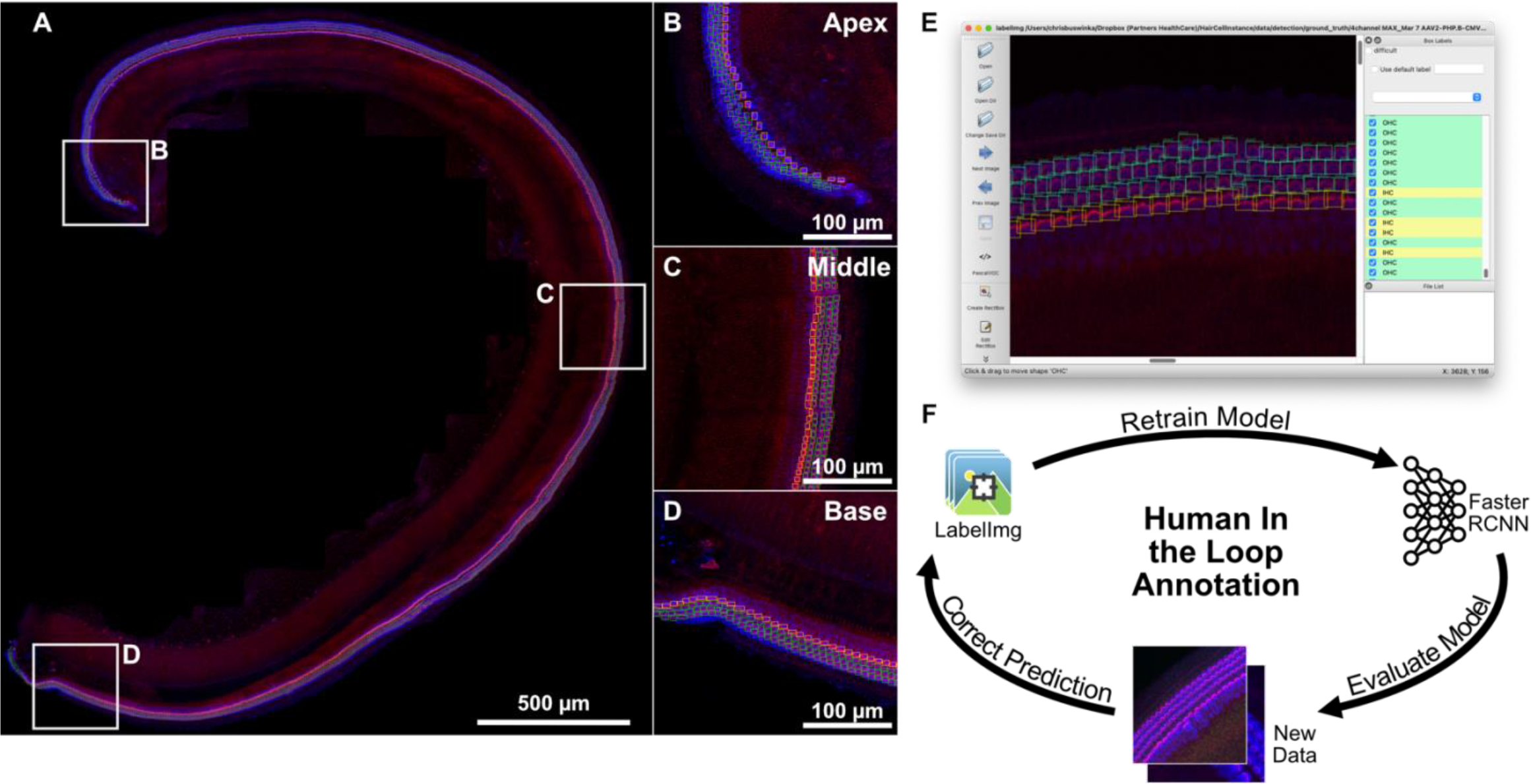
Cochlear hair cells annotation workflow using a ‘human-in-the-loop’ annotation paradigm. **(A)** An exemplar cochlea from our dataset with IHCs and OHCs annotated. Bounding boxes and classification labels were generated for all hair cells along the sensory epithelium. Representative regions in the **(B)** apex, **(C)** middle, and **(D)** base are shown as insets. Bounding boxes and labels were generated using labelImg **(E)**, an open-source object annotation software, with green boxes annotating OHCs and yellow boxes annotating IHCs. The annotation procedure was optimized using the human-in-the-loop paradigm **(F)**, where annotations were first generated by a preliminary neural network, then manually corrected and used to further train and improve the preliminary network, iteratively enabling more accurate candidate detections.

## MATERIALS and METHODS

### Sample Preparation and Imaging

#### Indzhykulian Laboratory

All experiments were carried out in compliance with ethical regulations and approved by the Animal Care Committee of Massachusetts Eye and Ear. Early postnatal (P3-5) murine cochleae were dissected as a single piece in Leibovitz’s L-15 culture medium (21083-027, Thermo Fisher Scientific) then fixed in 4% formaldehyde for 1 hour. In preparation for immunolabeling, samples were then permeabilized with 0.2% Triton-X for 30 minutes then blocked with 10% goat serum in calcium-free Hank’s Balanced Salt Solution (HBSS) for two hours. Hair cells were labeled with an anti-Myosin 7A primary antibody (#25-6790 Proteus Biosciences, 1:400) followed by application of goat anti-rabbit CF568 (Biotium) secondary antibody. Samples were additionally labeled with Phalloidin, allowing visualization of actin filaments (Biotium CF640R Phalloidin) and DAPI to visualize the nuclei. Stained samples were mounted on a slide using ProLong Diamond Antifade Mounting kit (P36965, Thermo Fisher Scientific).

Cochlear samples were imaged with a Leica SP8 confocal microscope (Leica Microsystems) using a 63×, 1.3 NA objective. Multiple confocal Z-stacks of 512×512 pixels in X and Y, with an effective pixel size of 288 nm, and a Z step of 1 µm were collected using the tiling function of the Leica LASX acquisition software to encompass whole cochleae. Stacks were maximum intensity Z-projected to form 2D images and annotated as outlined below.

#### Cunningham Laboratory

The images were generated as part of a previously reported study^16^. At the end of both cyclic drug administration protocols, mice were euthanized via carbon dioxide asphyxiation followed by decapitation. Cochleae were rapidly dissected and perfused with 4 % paraformaldehyde (PFA) at 4 °C through the round and oval windows and then post-fixed for 1 h at room temperature or overnight at 4 °C. Fixed tissue was decalcified in 0.5 M EDTA for 48 h at room temperature or for up to 96 h at 4 °C.

The cochleae of ears processed for whole mounts were microdissected in 1× phosphate-buffered saline (PBS) into 5 isolated turns to be immunostained and imaged. Tissue was incubated in blocking solution (5 % normal horse serum in 1× Phosphate Buffered Saline (PBS, Sigma-Aldrich) and Triton X-100 (Sigma-Aldrich; 1:300) for 1 h and then rinsed for 15 min in PBS. Cochlear turns were immunostained with antibodies to (1) myosin-VIIa (rabbit anti-myosin-VIIA; Proteus Biosciences, Ramona, CA; 1:200) and (2) C-terminal binding protein 2 (mouse anti-CtBP2; BD Biosciences; used at 1:200) with secondary antibodies coupled to Alexa Fluors 647 (Invitrogen; used at 1:200) and 568 (Invitrogen; used at 1:1000) respectively. In addition, cochlear turns were stained for actin using Alexa Fluor 488-conjugated phalloidin (Invitrogen; used at 1:50) for the equimolar cyclic drug administration experiment. Due to changes in microscope settings, this antibody was used at 1:1000 for some experiments. All antibodies were diluted in 1 % NHS and 30 % Triton X-100. Immunostained tissue was mounted on glass slides using Fluoromount-G (Southern Biotech).

A cochlear frequency map was created via a custom plug-in for ImageJ. Confocal z-stacks (step size of 0.2 μm) corresponding to representative apical (8 kHz), middle (16 kHz), and basal (44 kHz) regions from each ear were collected using an LSM 780 laser scanning confocal microscope (Carl Zeiss AG, Oberkochen, Germany) in a 1024 × 1024 pixel raster (135 μm^2^) using an oil-immersion objective (63×) of numerical aperture 1.4. IHCs and OHCs were counted in 75-μm-long stretches of the basilar membrane based on the nuclei labeled with C-terminus binding protein 2 (CtBP2). Images were then processed to generate maximum intensity projections and annotated as outlined below.

#### Ballesteros Laboratory

Early postnatal P1 or P6 murine cochleae were dissected as a single piece in Leibovitz’s L-15 culture medium (21083-027, Thermo Fisher Scientific) and then fixed in 4% PFA in Hank’s Balanced Salt Solution (HBSS) for 20 min at room temperature and under gentle shaking. Tissue was washed with PBS to remove PFA excess and permeabilized by incubation with 0.5% Triton-X in PBS for 30 minutes at room temperature. After permeabilization, tissue was washed three times with PBS and then blocked with 10% Normal goat serum (50062Z, Invitrogen). Hair cells were labeled with a 1:100 dilution of anti-Myosin 7A primary antibody (#25-6790 Proteus Biosciences) in goat serum overnight at 4C, followed by application of a 1:1000 dilution of Alexa Fluor-555 goat anti-rabbit (A32732, Invitrogen) and a 1:400 dilution of Alexa Fluor 488 phalloidin (A12379, Thermo Fisher Scientific) for one hour at room temperature. The tissue was then washed twice with PBS to remove the excess antibody. Stained samples were mounted on a slide using ProLong Diamond Antifade Mountant (P36965, Thermo Fisher Scientific). Animal care and experimental procedures were performed in accordance with the Guide for the Care and Use of Laboratory Animals and were approved by the Animal Care and Use Committee of the National Institute of Deafness and Other Communication Disorders (ASP1617).

Cochlear samples were imaged with an LSM980 confocal microscope equipment with an Airyscan 2 detector (Carl Zeiss), using an oil immersion alpha Plan-Apochromat 63X/1.4 Oil Corr M27 objective (Carl Zeiss) and Immersol 518F immersion media (ne=1.518 (30°C), Carl Zeiss). Multiple confocal Z-stacks of 512×512 pixels in X and Y, with a pixel size of 290 nm, were collected using the Zen (Carl Zeiss) acquisition software’s tiling function to encompass the whole organ of Corti epithelia.

#### Edge Laboratory

The images were generated as part of a previously reported study^17^. All experiments were carried out in compliance with ethical regulations and approved by the Animal Care Committee of Massachusetts Eye and Ear. Cochleae were harvested from *Ndp* knockout and wild-type mice in which alkaline phosphatase was inserted into exon 2 (Stock No. 011076 Jackson Laboratory) at the ages of postnatal day 3 (P3) and 2 months (2 mo), respectively. For whole mounts of P3 mice, sensory epithelia were fixed at room temperature for 15 minutes in 4% PFA in 0.1 M phosphate buffer (pH 7.4), then rinsed with PBS. For whole mounts of 2 mo mice, temporal bones were fixed at room temperature for 2 hours in 4% PFA in 0.1 M phosphate buffer (pH 7.4), then rinsed with PBS. After decalcification with 0.12 M EDTA (pH 7.0) and dehydration with 30% sucrose, surface preparation was performed. All specimens were incubated at room temperature for 30 minutes in 10% donkey serum with 0.2% Triton X-100 for blocking. Hair cells were labeled with an anti-Myosin 7A primary antibody (#25-6790 Proteus Biosciences, 1:500) followed by application of a donkey anti-rabbit AF488 secondary antibody (Invitrogen, 1:500). Samples were additionally labeled with Phalloidin, allowing visualization of actin filaments (Invitrogen AF647 Phalloidin, 1:100) and DAPI (Invitrogen,10 µg/ml) to visualize the nuclei. Stained samples were mounted on a slide using Mounting Medium with DAPI (H-1200 VECTASHIELD). Cochlear samples were imaged with a Leica SP8 confocal microscope (Leica Microsystems) using a 63×, 1.3 NA objective to generate confocal Z-stacks of 512×512 pixels in X and Y, with an effective pixel size of 360 nm. Stacks were maximum intensity Z-projected to form 2D images and annotated as outlined below.

#### Gnedeva Laboratory

Whole-mount P1-2 neonatal murine cochleae were dissected in 1X ice-cold PBS (Sigma) and fixed in 4% formaldehyde for 10 min. The organs were then blocked with 10% donkey serum in Tris Based Buffer supplemented with 0.1% Triton x100 (Sigma) and 0.01% Sodium Azide (Sigma). Hair cells were labeled with anti-Myosin 7A (Proteus Biosciences) or anti-Parvalbumin (Hudspeth Laboratory) primary antibodies and donkey anti-rabbit (Abcam) secondary antibodies. Phalloidin was used to visualize actin filaments (Sigma) and DAPI was used to visualize the nuclei. The samples were imaged with a Zeiss LSM 800 confocal microscope using a 20× objective. The images were collected at 1024×1024 pixels with a pixel size of 0.31um.

#### Landegger and Arnoldner Laboratories

Five-month-old pigs were euthanized at the end of cochlear implantation experiments in a prior study. Both inner ears of each animal were extracted with a hole saw and immediately stored in 4% formaldehyde (pH 7.4) for 24 hours. The contralateral, untreated side was decalcified for two months in 12% ethylenediaminetetraacetic solution (pH 7.4) at 37 °C and whole mounts were dissected in 1X phosphate-buffered saline. In preparation for immunolabeling, samples were permeabilized with 1% Triton-X and blocked simultaneously with 10% goat serum in 1× PBS for one hour. Hair cells were labeled with anti-Myosin 7A primary antibody (#25-6790 Proteus Biosciences, 1:200) followed by application of goat anti-rabbit 568 secondary antibody (Invitrogen, #A11011, 1:400). Samples were additionally labeled with Phalloidin for visualization of actin filaments (Alexa Fluor 488, Invitrogen #A12379, 1:500) and DAPI to visualize the nuclei (1:1000). Stained samples were mounted on a side using ProLong Diamond Antifade Mounting kit (#P36930, Thermo Fisher Scientific). All experiments were approved by the Animal Care Committee of the Medical University of Vienna and by the local animal welfare committee and the Austrian Federal Ministry of Education, Science and Research (BMBWF-2020–0.272.252). Cochlear samples were imaged with a Nikon Ti Eclipse confocal microscope (Nikon, Tokyo, Japan) and a 60× 1.4NA oil immersion objective. Multiple confocal Z-stacks of 1024×1024 pixels in X and Y (207nm/pix) were generated with a distance of 2.5 µm between each image scan in Z. All stacks were Z-projected at their maximum intensity to form 2D images and annotated as outlined below.

#### Liberman Laboratory

The images were generated as part of a previously reported study^8^. Temporal bones were extracted at autopsy with a bone-plugging tool soon after death and immediately immersed in buffered 10% formalin after opening the round and oval windows. All procedures were approved by the Human Studies Committee of the Massachusetts Eye and Ear Infirmary. After post-fixation (4°C) for at least 6 days, the bone plug containing the cochlea was drilled to remove as much of the petrous bone as possible, and then immersed in EDTA at room temperature for ∼27 days. The cochlea was then microdissected into 8 – 9 pieces, each containing the osseous spiral lamina and the attached organ of Corti. Then, each piece underwent a freeze/thaw step in 30% sucrose for permeabilization, followed by 1 hr at room temperature in a blocking buffer (PBS with 5% normal horse serum and 0.3 -1% Triton X-100). Tissue was then incubated overnight at 37 °C with the following primary antibodies (plus 0.3 – 1% TritonX): 1) rabbit anti-Myosin VI and/or VIIa (Proteus Biosciences #25-6791 and 25-6790, respectively) at 1:100 to count hair cells, 2) mouse (IgG1) anti-CtBP2 (C-terminal Binding Protein; BD Biosciences #612044) at 1:200, 3) chicken anti-neurofilament (Chemicon #AB5539) at 1:1000, and 4) goat anti-ChAT (choline acetyltransferase; Millipore #AB144P) at 1:100. Primary incubations were followed by 2 sequential 60-min incubations at 37°C in species-appropriate secondary antibodies (coupled to Alexa Fluor dyes) with 0.3 -1% Triton-X. Confocal z-stacks of the inner and outer hair cells were acquired at 14 equally spaced locations along the spiral at 240 nm/pixel in x and y (digital zoom 0.75) and with 0.33 µm z-spacing on a Leica SP8 using a 63× glycerol objective (1.3 N.A.).

#### Manor Laboratory

Samples were prepared as described in the Indzhykulian Laboratory procedures above, then imaged with two microscope systems. For Airyscan imaging, cochlear samples were imaged with a Zeiss LSM 880 Rear Port Laser Scanning Confocal and Airyscan FAST microscope using a 63×, 1.4 NA objective. Multiple confocal Z-stacks of 1792 × 1792 pixels in X and Y, with an effective pixel size of 43 nm, and a Z step of 0.5 mm were collected using the tiling function of the Zeiss Zen Black 2.3 acquisition software to encompass representative basal, middle, and apical cochlear regions. For spinning disk imaging, cochlear samples were imaged with a Zeiss CSU Spinning Disk Confocal microscope equipped with a Yokogawa spinning disk scan head and an EM-CCD camera using a 40×, 1.3 oil DIC objective. Multiple confocal Z-stacks of 11180×8944 pixels in X and Y, with an effective pixel size of 229 nm, and a Z step of 0.5 mm were collected using the tiling function of the Zeiss Zen Blue 2.3 software to encompass whole cochleae.

#### Ricci Laboratory

The images were generated as part of a previously reported study^18^. All animal studies were conducted in strict accordance with the protocols approved by the Institutional Animal Care and Use Committee at Stanford University (APLAC-14345). We utilized C57BL/6 mice (Charles River: 026) of both sexes, aged between 4 and 8 weeks. Gentamicin (180 mg/kg) was systematically administered through intraperitoneal (IP) injection. Thirty minutes later, furosemide (100 mg/kg) was also injected to expedite gentamicin-induced ototoxicity. These injections were repeated daily for a period of 10 days. The cochleae were harvested from the animals and fixed with 4% PFA/PBS (15710, EMS). To decalcify the tissue, 0.5M EDTA (E177, VWR) was applied for six hours. We dissected the whole-mount organ of Corti and removed the tectorial membrane. Subsequently, the tissue was permeabilized for 30 minutes in 0.5% Triton X-100/PBS (BP151, Fisher Scientific) and then blocked for two hours in 5% BSA/PBS (BP1600, Fisher Scientific, Fair Lawn, NJ) at room temperature. For staining, we employed primary antibodies against MyosinVIIa (hair cell marker; 1:500; MYO7A 138-1, DSHB), which were incubated overnight at 4°C and then for an additional two hours at 37°C in the same blocking buffer. Following three washes with the blocking buffer, the tissues were incubated with secondary antibodies conjugated with Alexa Fluor 546 (1:500; A10036, Life Technologies) and Alexa Fluor 647-conjugated phalloidin (F-actin marker; 1:200; A22287; Invitrogen) for an hour at room temperature. After five washes with PBS, the tissues were mounted on a glass slide using ProLong™ Diamond Antifade Mountant with DAPI (P36962, Invitrogen) and stored in the dark at 4°C. Z-stack images of cochlear tissues were acquired using a confocal microscope (LSM700, Zeiss, Jena, Germany) equipped with a 40× objective (NA 1.3). Each individual image comprises a resolution of 1660 × 1660 pixels (X × Y) with a pixel size of 96 nm. The images were collected at intervals of 1 µm along the Z-axis. Z-stack projections using average intensity were generated using ImageJ software.

#### Richardson and Kros Laboratories

The images were generated as part of a previously reported study^19^ assaying hair cell survival in response to aminoglycosides and putative protective compounds from the Life Chemicals Diversity Set. Briefly, P2 mouse cochlear cultures were prepared from wild-type CD-1 mice of either sex. Organs of Corti were plated onto collagen-coated (Corning, 354236) coverslips in 93% DMEM-F12, 7% FBS, and 10 μg/mL ampicillin, and maintained for 24 hours at 37°C in Maximow slide assemblies. Mouse colonies were maintained at the University of Sussex and kept in full accordance with United Kingdom Home Office regulations.

Following 24 hours of incubation, coverslips with adherent cochleae were placed in 35 mm Petri dishes (Greiner Bio-One, 627161), and 1 mL culture media (98.8% DMEM/F12, 1.2% FBS) containing either vehicle (0.5% DMSO), 5 μM gentamicin with 0.5% DMSO, or 5 μM gentamicin with 50 μm test compound UoS-7692. After 48 hours, cultures were washed in PBS, fixed at room temperature for 1 hour in 3.7% formaldehyde (v/v) (MilliporeSigma, F1635) in 0.1 M sodium phosphate pH 7.4. For labelling cultures were incubated at 4°C overnight in PBS containing 10% horse serum and 0.1% Triton X-100 with 1:200 Texas red phalloidin (Invitrogen, Thermo Fisher Scientific, T7471) and 1:1000 rabbit anti-myosin VIIa (Proteus Bio-Sciences, 25-6790), followed by 4 hours at room temperature in 1:500 Alexa Fluor 488 goat anti-rabbit secondary antibody (Invitrogen, Thermo Fisher Scientific, A-11034). Cultures were mounted in Vectashield (Vector Laboratories, H-1000) and imaged in a mid-basal region, approximately 20% along the length of the cochlea from the basal end, using a 40× 0.75 NA objective on a Zeiss Axioplan2 upright microscope with a Spot RT Slider camera with an effective pixel size of 183 nm. In order to facilitate quantification, images were obtained from multiple focal planes and merged into one image per channel in which all cells were in focus using Adobe Photoshop Creative Cloud.

#### Rutherford and Deans Laboratories

All procedures were approved by the ASC at Washington University in St. Louis and IACUC at the University of Utah in Salt Lake City. Samples were prepared as previously described (Ghimire and Deans, 2019; Kim et al., 2019; Hu et al., 2020; Walia et al., 2021; Boero et al., 2021; Rutherford et al., 2023). Briefly, mice were euthanized by CO_2_ inhalation and perfused through the heart with fixative solution (4% PFA in sodium phosphate buffer). Temporal bones were isolated in PBS. A hole was made through the bone near the cochlear apex, the round and oval windows were punctured, and the temporal bone was bath fixed in fixative solution for one hour at 4 °C. The samples were decalcified in 10% EDTA for 3 hours. Whole-mount preparations containing organ of Corti and spiral ganglion were isolated from the cochlea in three pieces containing the apical, middle, or basal thirds. After incubation in blocking buffer conaining 10% donkey serum and 0.2% Triton-X at room temperature for 2 hours, antibodies were applied in blocking buffer at 4 °C overnight: (Myosin7a rabbit, Proteus Biosciences; Alexa Fluor 555 Phalloidin). The Myosin7a antibody was then labelled with AlexaFluor 488 for 1 hour at room temperature. Samples were mounted on slides in Mowiol mounting medium and imaged on a Zeiss LSM700 with a z-step of 0.5 micrometers and pixel size of 100 nm in x and y with a 63× 1.4NA oil objective lens (Carl Zeiss). Image stacks were maximum intensity Z-projected to form 2D images and annotated as described below.

#### Stepanyan and Basch Laboratories

Samples were prepared as described in the Indzhykulian Laboratory procedures above, then imaged on Zeiss Structure Illumination Microscope system with 20× 0.8 NA and 40× 1.3 NA objective lenses. Zen 2.3 pro software was used for image acquisition with an Axiocam 702 mono camera (1920×1216 pixels). Images were collected as a Z-stack with a 1 μm step and 294 nm (20× lens) or 147 nm (40× lens) pixel size. Image stacks were then maximum intensity Z-projected to form 2D images and annotated as described below.

#### Shrestha Laboratory

All experiments were carried out in compliance with regulations and procedures approved by the Institutional Animal Care and Use Committee (IACUC) of Massachusetts Eye and Ear. Postnatal day 18 (P18) mice of both sexes were euthanized by administering an overdose of pentobarbital sodium (Fatal Plus, 150 mg/kg). Temporal bones were quickly removed and immersed for 1 hour in fresh 4% PFA in 1× PBS. They were then decalcified in 120 mM EDTA at 4°C for 48 hours. Cochleae were dissected out and stored in 1× PBS at 4 °C until further use 24-72 hrs later. Immunostaining was performed on these whole mount preparations as follows: they were permeabilized and blocked for 1 hr in 1x PBS with 1% Triton X-100 (PBS-Tx) and 5% normal donkey serum (NDS). Tissues were then incubated overnight at 4°C with rabbit anti-Parvalbumin (1:250, Swant #PV27a) in 0.3% PBS-Tx and 1% NDS. After tissues were rinsed in PBS-Tx for 45 mins, they were incubated for 2 hours at room temperature with Alexa Fluor 555 donkey anti-rabbit (1:500) and Alexa Fluor 488 Phalloidin (1:500, Invitrogen #A12379) in 0.3% PBS-Tx and 1% NDS. After rinsing again for 45 minutes in PBS-Tx, all tissues were mounted in Fluoromount-G (Electron Microscopy Sciences, #17984-25). Images were acquired from the middle turn at 90 nm/pixel XY resolution (1024 × 1024 pixels) and 0.33 μm optical sectioning in the Z dimension using a Leica SP8 confocal microscope fitted with a 63× 1.3 NA objective (digital zoom factor 2). Maximum intensity projections of these image stacks were created in Fiji. The resulting images were then annotated as outlined below.

#### Tarchini Laboratory

The images were generated as part of a previously reported study^20^. Animals were maintained under standard housing and all animal work was reviewed for compliance and approved by the Animal Care and Use Committee of The Jackson Laboratory. Briefly, temporal bones of postnatal day 21-22 mice were isolated, and the cochlea punctured at the apex before immersion fixation in 4% PFA for 1h at 4°C. Temporal bones were then incubated overnight at room temperature in 0.11M EDTA for decalcification before exposing the auditory epithelium. Samples were blocked and permeabilized in PBS with 0.5% Triton-X100 and bovine serum albumin (1%) for at least 1h at room temperature. Primary and secondary antibodies were each incubated overnight at 4°C in PBS. After each antibody incubation, samples were washed 3 times with PBS + 0.05% Triton-X100 before a final post-fixation in PFA 4% for 1h at room temperature. Antibodies used were rabbit MYO6 (Proteus; 25-6791) and mouse MYO7 (DSHB; MYO7A 138-1). Secondary antibodies were raised in donkey and coupled to Alexa Fluor (AF) 555 or 647 (ThermoFisher Scientific; anti-rabbit AF555, A-31572; anti-mouse AF555, A-31570; anti-mouse AF647, A-31571). We used fluorescent conjugated phalloidin to reveal F-Actin (ThermoFisher Scientific: AF488, A12379). Samples were then mounted flat on a microscopy slide (Denville M1021) directly under a 18×18mm #1.5 cover glass (VWR 48366-045), using Mowiol (10% w/v) (Calbiochem/MilliporeSigma 4759041) as mounting medium. Images were captured with a LSM800 line scanning confocal microscope using the Zen2.3 or Zen 2.6 software, the Airyscan detector in regular confocal mode, and a 63x 1.4 NA oil objective lens (Carl Zeiss AG). Confocal stacks were taken at the cochlear base, with a z-step of 0.1μm, a pixel size of 68nm, and fields of 1495 × 1495 pixels. Stacks were maximum intensity Z-projected to form 2D images and annotated as outlined below.

#### Vélez Laboratory

All animal procedures were approved by the University of Kentucky Animal Care and Use Committee (protocol 2020-3535). Organ of Corti explants were isolated from C57BL/6 mice at postnatal day 4 in cold Leibovitz’s L-15 medium (cat. # 21083027, Gibco) and held in place by two glass fibers glued to the bottom of a Petri dish. Explants were cultured in high-glucose DMEM (12430062, Gibco) supplemented with 7% FBS (16140071, Gibco) and 10 mg/L ampicillin (171254, Calbiochem) at 37°C and 5% CO2, for 24 h in the presence of 62.5 µM gentamicin (G1397, Sigma-Aldrich) or in control conditions. Next, explants were fixed in 4% PFA (15710, Electron Microscopy Sciences) for 24h at 4°C. Samples were then permeabilized with 0.5% Triton X-100 (22142, Electron Microscopy Sciences) for 1h and blocked with 10% normal goat serum (10000C, Invitrogen) and 0.25% Triton X-100 for 1 h, at room temperature. Hair cells were labeled with rabbit polyclonal anti-myosin VIIa primary antibody (25-6790, Proteus Biosciences, concentration 1:100) for 24 h at 4°C followed by labeling with goat anti-rabbit Alexa fluor 488 (A11034, Invitrogen, 1:1000) as a secondary antibody for 3 h at room temperature. Samples were counterstained with 1 unit of rhodamine phalloidin for 30 min at room temperature (R415, Invitrogen), and mounted in ProLong Diamond antifade medium (P36961, Invitrogen). Imaging was performed with a Leica SP8 upright confocal microscope equipped with a Leica HCX PL APO 100X 1.44 NA objective lens. Confocal z-stacks were taken with a voxel size of 114 nm in X and Y, and 500 nm in Z.

#### Walters Laboratory

All use of animals for these studies was approved by the Institutional Animal Care and Use Committee at the University of Mississippi Medical Center. Guinea pigs (*Cavia albino*) around the age of 15-17 weeks old were euthanized by deep anesthesia with isoflurane followed by a 200 mg/kg injection of Fatal Plus (Vortech Pharmaceuticals). After confirmatory decapitation, the temporal bones were removed, trepanned, and perfused with a solution of 4% PFA in PBS and then immersed in 4% PFA in PBS for 24 hours at 4 °C. The temporal bones were then decalcified in 0.25M ethylenediaminetetraacetic acid (EDTA) for 5 days on a paddle rotator at room temperature. The decalcified bony labyrinths were then dissected to remove the sensory epithelia which were stored in PBS until ready for immunostaining. Free floating samples were washed several times in PBS, blocked for 1 hour in 10% normal goat serum, 1% bovine serum albumin, and 1% Triton-X in PBS, then immersed in primary antibodies in blocking buffer that was diluted by half in PBS. The antibodies used were Rabbit anti MYO7A (Proteus Biosciences, #25-6790) @1:200, and Mouse anti-BRN3C (Santa Cruz Biotechnology, #sc-81980) @ 1:250. The samples were next washed 3 × 10 min. in PBS, then immersed in diluted blocking buffer containing Alexa-Fluor-conjugated secondary antibodies (Goat anti-Rabbit and Goat anti-Mouse IgG1, Invitrogen / Thermo-Fisher), each at 1:800 dilution, for 3 hours at room temperature. Cell nuclei in all samples were counterstained with Hoechst33258 at 1:1500 in PBS for 30 minutes and a subset of samples were simultaneously labeled by the addition of methanol reconstituted, AlexaFluor conjugated, phalloidin (Invitrogen/ThermoFisher) at 1:200 into the diluted Hoechst solution. Samples were then washed 3 × 5 minutes in PBS and mounted to slides using FluoroGel with DABCO (Electron Microscopy Sciences, #17985) and 1.5 thickness coverglass (Fisherbrand, Fisher Scientific). Z-stack images were then acquired under a 20× objective (0.8 NA) using a Zeiss LSM880 point-scanning confocal microscope. Large fields of view were collected using an XY resolution of 2048×2048 and a Z-slice thickness around 5 µm and 188 nm pixel size. Images were then processed to generate maximum intensity projections and annotated as outlined below.

#### Zhao Laboratory

Animal experiments were carried out in accordance with the National Institutes of Health Guide and were approved by the Institutional Animal Care and Use Committee of Indiana University School of Medicine. Cochlear explants dissected from P3-P4 mice were cultured overnight in DMEM/F12 media at 37°C in a 5% CO_2_ humidified atmosphere. Then, cochlear explants were exposed to saline, cisplatin, or other chemical solutions for two days. Samples were fixed in a fixative containing 4% PFA in HBSS for 20 min. After washing with HBSS, the tectorial membrane was removed. After blocking with HBSS containing 5% bovine serum albumin (BSA) and 0.5% Triton X-100 at room temperature for 20 minutes, samples were incubated with primary antibodies overnight at 4°C. Samples were washed in HBSS and incubated with secondary antibodies for 2 hours at room temperature. Then, tissues were mounted in ProLong® Antifade Reagents (ThermoFisher Sci). Stacked images were captured by a DM6 FS automated deconvolution microscope (Leica) using a 20× objective (HC PL FLUOTAR 20x/0.55; 214 nm/pixel). Antibodies used in this study were anti-βII spectrin (1:200, cat# sc-136074, Santa Cruz) and Alexa Fluor 488 goat anti-mouse (1:2,000, cat# A11017, Life technologies Corporation).

### Annotation Procedure

To qualify for annotation, cells were required to have a clear and definable central feature, either a well stained cuticular plate or a stereocilia bundle, connected to a labeled cell body of the expected shape and size for that tissue type/preparation. Most of the images within this collection were of hair cells with relatively normal appearance, with some instances of missing hair cells. Many of the annotated images are Z-projections, often with the cytosol of one cell occluding another rendering the cytosol as an unreliable feature for bounding box detection. Thus, each hair cell annotation box bounds the cuticular plate enclosing the stereocilia bundle and is assigned a class label of cell type (IHC or OHC).

Bounding box annotations were created and corrected in the labelImg open-source software^21^ and other scripts. The annotations were generated using a “human-in-the-loop”^22^ paradigm as outlined in **Figure 1**. Initial candidate annotations were generated using a neural network, which were then audited and refined to ensure every cell classification label was correct, and that bounding boxes tightly enclose the cell’s stereocilia bundle without including any additional features. Since the images within the dataset were annotated by several annotators, each image was then reviewed by a single lead annotator to ensure accuracy and minimize inter-annotator biases. Data annotations were saved as a separate xml file in the coco format with an identical filename to the associated image^23^. Although uncommon, some images within the dataset were collected following application of certain insults and may include hair cells with damaged stereocilia bundles.

### DATA RECORDS

The dataset presented here is hosted with Zenodo^24^ data repository and is subdivided by type of microscopy, type of treatment, species and research group of origin. Collectively we have annotated a total of **94**,**095** hair cells, over 553 images from over 20 research groups^25^. The dataset presents examples of auditory hair cells of four species: Mouse, Guinea Pig, Pig and Human, imaged at differing pixel sizes, magnifications, using point scanning confocal, spinning disk confocal, structured illumination, or widefield fluorescence microscopy techniques. Depending on origin, individual images may either contain the entire cochlea or smaller sub-regions. A detailed summary of the dataset, along with the relevant parameters is presented in **Table 1**. The materials and methods used by each group are reported within the methods section of this manuscript.

Details of sample preparation, stain, and resolution are embedded in its annotation metadata. We present each maximum intensity projection image in the dataset in the TIFF-format, along with associated ground truth detection annotations in XML. For ease of use, we have created a small python library, which can automatically download and parse the dataset into a code-accessible format. As many of our manually annotated images are of full cochleae, we included exemplar functionality to selectively sample each image when training, allowing precise control to correct for class imbalance. In this way, an underrepresented image type may be showed to the model more often, improving overall generalizability. Finally, we present an exemplar data augmentation procedure for our dataset which can aid in training accurate models in the future. Detailed descriptions of these procedures were previously reported^10^ and briefly summarized within the methods section.

### TECHNICAL VALIDATION

To ensure the accuracy of our dataset, we employed an iterative process whereby a set of expert annotators correct predicted candidate annotations from an existing deep-neural network. The process is as follows: 1) the detection neural network was trained on a preliminary subset of images with manual annotations, 2) the resulting network was evaluated on new images to generate candidate annotations, 3) these candidate annotations were inspected and corrected (as needed) by at minimum two trained annotators followed by 4) periodical re-training the detection network using newly generated pool of ground truth annotations and repeating the process. Priority was given to manually corrected poorly performing examples at each iterative stage, thereby allowing the network to direct the annotation efforts toward generalizability. Final candidate annotations were always corrected and approved by the lead annotator to eliminate any possible systematic differences in annotation style between annotators and ensuring high quality. In some cases, images contain examples of atypical cells which are challenging to annotate. In these cases, researchers from the laboratory of origin were consulted on annotations. We have further validated the utility of this dataset in a previously reported study^10^ and show that this dataset on its own is sufficient to generate accurate deep-learning detection models.

### USAGE NOTES

Our dataset contains images which represent vastly different spatial areas. Many of these images are too large to pass through a deep neural network. Therefore, we recommend sampling equal sized regions of each image a pre-determined number of times. Class and frequency imbalance issues can therefore be mitigated by choosing the sampling factor for each image, or each group of images based on the species, animal’s age, type of insult, the laboratory of origin, etc. Furthermore, as the cochlea is a spiraled organ, much of the area in some of our larger images contain no hair cells (center of the spiral), therefore we recommend centering the sampled region by an existing cell annotation.

Often, a plethora of data is not enough to prevent overtraining in deep neural networks. We therefore recommend extensive data augmentation when using this dataset for training. Of critical importance are affine transformations, which randomly rotate, shear, and scale the data. This can be challenging when working with bounding boxes, as they may increase in area when rotated. Thus, we recommend shrinking rotated boxes slightly when rotated. In general, augmentation of training data makes a resulting model invariant to the augmentations. Therefore, we recommend the following augmentations to improve model generalizability: 1) shuffle the order of fluorescence channels (switching colors) on an image to simulate changes in image structure (i.e., MYO7A is not always green, phalloidin is not always red, etc.), 2) adjust the brightness and contrast of each channel separately to simulate weak or strong staining, 3) add random noise to simulate low signal-to-noise ratio imaging, and 4) blur the image to reduce dependency of the model to proper focus or high resolution. A detailed description of our recommended augmentation procedure was previously reported^10^ and successfully implemented in the associated data augmentation pipeline.

## DATA AVAILABILITY

All imaging data along with the annotations can be found at: https://zenodo.org/record/7937969^25^.

## CODE AVAILABILITY

All associated code for downloading, loading, and preprocessing this dataset may be found at: https://github.com/indzhykulianlab/hcat-data.

## ACKNOWLEDGEMENTS

We would like to thank Haobing Wang, MS (Mass Eye and Ear Light Microscopy Imaging Core Facility) for their assistance in this project.

## AUTHOR CONTRIBUTIONS

Christopher J. Buswinka, Conceptualization, Methodology, Software, Validation, Investigation, Data Curation, Visualization, Supervision, Writing – Original Draft.

David B. Rosenberg, Data Curation, Writing – Review & Editing.

Rubina G. Simikyan, Data Curation, Writing – Review & Editing.

Richard T. Osgood, Data Curation, Validation, Writing – Review & Editing Katharine Fernandez, Data Acquisition, Writing-Review & Editing.

Yushi Hayashi, Data Acquisition, Writing-Review & Editing.

Hidetomi Nitta, Data Curation, Writing – Review & Editing.

Leslie W. Liberman Data Acquisition, Writing-Review & Editing.

Emily Nguyen, Data Curation, Writing-Review & Editing.

Erdem Yildiz, Data Acquisition, Writing-Review & Editing. Jinkyung Kim, Data Acquisition, Writing-Review & Editing.

Amandine Jarysta, Data Acquisition, Writing-Review & Editing.

Justine Renauld, Data Acquisition, Writing-Review & Editing.

Ella Wesson, Data Acquisition, Writing-Review & Editing.

Punam Thapa, Data Acquisition, Writing-Review & Editing Pierrick Bordiga, Data Acquisition, Writing-Review & Editing.

Noah McMurtry, Data Acquisition, Writing-Review & Editing.

Juan Llamas, Data Acquisition, Writing-Review & Editing.

Siân R. Kitcher, Data Acquisition, Writing-Review & Editing.

Ana I. López-Porras, Data Acquisition, Writing-Review & Editing.

Runjia Cui, Data Acquisition, Writing-Review & Editing.

Ghazaleh Behnammanesh, Data Acquisition, Writing-Review & Editing.

Jonathan E. Bird, Data Acquisition, Writing-Review & Editing.

Angela Ballesteros, Data Acquisition, Writing-Review & Editing.

A. Catalina Vélez-Ortega, Data Acquisition, Writing-Review & Editing.

Albert SB Edge, Data Acquisition, Writing-Review & Editing.

Michael Deans, Data Acquisition, Writing-Review & Editing.

Ksenia Gnedeva, Data Acquisition, Writing-Review & Editing.

Brikha R. Shrestha, Data Acquisition, Writing-Review & Editing.

Uri Manor, Data Acquisition, Writing-Review & Editing.

Bo Zhao, Data Acquisition, Writing-Review & Editing.

Anthony J. Ricci, Data Acquisition, Writing-Review & Editing.

Basile Tarchini, Data Acquisition, Writing-Review & Editing.

Martin Basch, Data Acquisition, Writing-Review & Editing.

Ruben S. Stepaynan, Data Acquisition, Writing-Review & Editing.

Lukas D. Landegger, Data Acquisition, Writing-Review & Editing.

Mark Rutherford, Data Acquisition, Writing-Review & Editing.

M. Charles Liberman, Data Acquisition, Writing-Review & Editing.

Bradley J. Walters, Data Acquisition, Writing-Review & Editing.

Corne Kros, Data Acquisition, Writing-Review & Editing.

Guy P. Richardson, Data Acquisition, Writing-Review & Editing.

Lisa L. Cunningham, Data Acquisition, Writing-Review & Editing.

Artur A. Indzhykulian, Conceptualization, Project administration, Supervision, Writing – Review & Editing, Resources, Funding acquisition.

